# Induction of hierarchy and time through one-dimensional probability space with certain topologies

**DOI:** 10.1101/780882

**Authors:** Shun Adachi

## Abstract

In a previous study, the authors utilized a single dimensional operationalization of species density that at least partially demonstrated dynamic system behavior. For completeness, a theory needs to be developed related to homology/cohomology, induction of the time dimension, and system hierarchies. The topological nature of the system is carefully examined and for testing purposes, species density data for a wild Dictyostelia community data are used in conjunction with data derived from liquid-chromatography mass spectrometry of proteins. Utilizing a Clifford algebra, a congruent zeta function, and a Weierstraß *℘* function in conjunction with a type VI Painlevé equation, we confirmed the induction of hierarchy and time through one-dimensional probability space with certain topologies. This process also served to provide information concerning interactions in the model. The previously developed “small *s*” metric can characterize dynamical system hierarchy and interactions, using only abundance data along time development.

## 1 Introduction

In a previous study, the authors developed a system whereby a static set of species density information can be utilized to predict dynamics therein by extracting probabilistic information [1]. We developed a new complex system measure, “small *s*”, related to a probability space. When *N*_*k*_ is the individual density for the *k*-th ranked species and is approximated by a logarithmic distribution with parameters *a*, *b* with respect to the ranks of the values of individual densities,

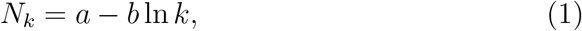

and

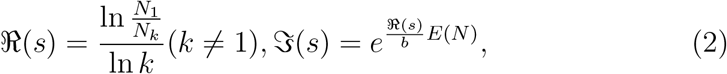

where *E*(*N*) is averaged species density. For *k* = 1, 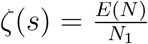 for species, where ζ(*s*) is a Riemann zeta function. Therefore, it appears doubtful why single-dimensional information (*N*_*k*_), with a topology labelled by rank *k*, can induce a 3-dimensional system (*a*, *b*, ln *k*, regarding *N_k_* as free energy, the others as internal energy or enthalpy, temperature, and entropy, respectively) of an individual density, accompanied with an even additional time dimension. To explain this, first of all, we set a 1-dimensional *C*^∞^ manifold with a topology as 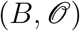 with *s* ∈ *B*. Inspired by the Bethe ansatz (e.g. [2]), we set three different topologies isomorphic to Δ, ℂ, 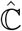 for further clarification of our model. These topologies naturally invest a cohomology, time dimension, and hierarchy to the system. Furthermore, we are able to define a proper topology independently from moduli of measurements with individual numbers and a Galois action dependent on moduli of it in an evolutionary system with hierarchy by Galois extension, such as biological systems in this case. For application to biological hierarchies, this model is tested using protein abundance data derived from liquid-chromatography mass spectrometry (LC/MS) of HEK-293 cells and species density data from a wild Dictyostelia community. Finally, we sought to evaluate interactions of the constituents of biological systems by invoking a Weierstraß *℘* function to estimate the strength of homo- and hetero-interactions. These results serve to further justify our “small *s*” metric to decipher system dynamics of interest. For example, adapted, non-adapted (neutral), and disadapted (repressed) proteins can be classified by expansion of the model using a Clifford algebra. Furthermore, utilizing a congruent zeta function elucidates the contribution to adaptive/disadaptive situations from each hierarchy.

## 2 Field Research & Experiments

### 2.1 Field Research

Data concerning the number of individuals in each species were obtained from natural (nonlaboratory) environments. The sampling is described in [3]. Field experiments were approved by the Ministry of the Environment, Ministry of Agriculture, Forestry and Fisheries, Shizuoka Prefecture and Washidu Shrine (all in Japan). The approval Nos. are 23Ikan24, 24Ikan72-32, and 24Ikan72-57. Soil samples were obtained from two point quadrats in the Washidu region of Izu in Japan. The number of individual cellular slime molds per gram of soil was determined by counting the number of plaques cultivated from soil samples. Species were identified by morphology and the DNA sequences of 18S rRNA genes. Samples were obtained monthly from May 2012 to January 2013 inclusive. Relevant calculations were performed using Microsoft Excel 16.16.13 and SageMath 8.8.

In more detail, sampling occurred using two 100 m^2^ quadrats in Washidu (35°3′33′′N, 138°53′46′′E; 35°3′45′′N, 138°53′32′′E). Within each 100 m^2^ quadrat, nine sample points were established at 5 m intervals. From each sampling point, 25 g of soil was collected. Cellular slime molds were isolated from these samples as follows. First, one sample from each site was added to 25 ml of sterile water, resuspended, and then filtrated with sterile gauze. Next, 100 *μ*l of each sample solution was mixed with 100 *μ*l HL5 culture medium containing *Klebsiella aerogenes* and spread on KK2 agar. After two days of storage in an incubator at 22 °C, the number of plaques on each agar plate was enumerated and recorded. Note that the number of plaques corresponds to the total number of living cells at any possible stage of the life cycle. That is, the niche considered here is the set of propagable individuals of Dictyostelia; these are not arranged in any hierarchy or by stage in the life cycle. Also, note that we did not examine the age or size structure of organisms, since most of these were unicellular microbes. Mature fruiting bodies consisting of cells from a single species were collected along with information regarding the number of plaques in the regions in which each fruiting body was found. Finally, spores were used to inoculate either KK2 for purification or SM/5 for expansion. All analyses were performed within two weeks from the time of collection. The isolated species were identified based on 18S rRNA (SSU) sequences, which were amplified and sequenced using PCR/sequencing primers, as described in [4] and the SILVA database (http://www.arb-silva.de/). The recipes for the media are described at http://dictybase.org/techniques/media/media.html.

### 2.2 Experiments

#### 2.2.1 Cell culture

A human HEK-293 cell line from an embryonic kidney was purchased from RIKEN (Japan). The sampling is described in [5]. The original cultures were frozen on either March 18, 2013 (3-year storage) or March 5, 2014 (2-year storage). They were subsequently used in experiments between February and June 2016. The strain was cultured in Modified Eagle’s Medium (MEM) + 10% fatal bovine serum (FBS) + 0.1 mM nonessential amino acid (NEAA) at 37 °C with 5% CO_2_. Subculturing was performed in 0.25% trypsin and prior to the experiment, the original cells from RIKEN were frozen following the standard protocol provided by RIKEN: in culture medium with 10% dimethyl sulfoxide (DMSO), they were cooled until reaching 4 °C at −2° C/min, held at that temperature for 10 min, then cooled until reaching −30° C at −1 °C/min in order to freeze, held at that temperature for 10 min, then cooled again until reaching −80 °C at −5 °C/min, and finally held at that temperature overnight. The next day, they were transferred to storage in liquid nitrogen.

#### 2.2.2 Protein experiments

The HEK-293 proteins were extracted using the standard protocol for the RIPA buffer (NACALAI TESQUE, INC., Kyoto, Japan). The sampling is described in [5]. Approximately 10^6^ harvested cells were washed once in Krebs-Ringer-Buffer (KRB; 154 mM NaCl, 5.6 mM KCl, 5.5 mM glucose, 20.1 mM HEPES pH 7.4, 25 mM NaHCO3). They were resuspended in 30 *μ*l of RIPA buffer, passed in and out through 21G needles for destruction, and incubated on ice for 1 h. They were then centrifuged at 10,000 g for 10 min at 4 °C, followed by collection of the supernatants. The proteins were quantified using a Micro BCA Protein Assay Kit (Thermo Fisher Scientific, Waltham, U.S.A.) and further processing was performed using XL-Tryp Kit Direct Digestion (APRO SCIENCE, Naruto, Japan). The samples were solidified in acrylamide gel, washed twice in ultrapure water, then washed three times in dehydration solution, and finally dried. The samples were then processed using an In-Gel R-CAM Kit (APRO SCIENCE, Naruto, Japan). The samples were reduced for 2 h at 37 °C, alkylated for 30 min at room temperature, washed five times with ultrapure water, washed twice with destaining solution, and then dried. The resultant samples were trypsinized overnight at 35 °C. The next day, the dissolved digested peptides were collected by ZipTipC18 (Merck Millipore, Corp., Billerica, U.S.A.). The tips were dampened twice with acetonitrile and equilibrated twice with 0.1% trifluoroacetic acid. The peptides were collected by ~20 cycles of aspiration and dispensing, washed twice with 0.1% trifluoroacetic acid, and eluted by 0.1% trifluoroacetic acid /50% acetonitrile with aspiration and dispensing five times × three tips followed by vacuum drying. The final samples were stored at −20 °C. Before undertaking LC/MS, they were resuspended in 0.1% formic acid, and the amounts were quantified by Pierce Quantitative Colorimetric Peptide Assay (Thermo Fisher Scientific, Waltham, U.S.A.). This protocol is published at http://dx.doi.org/10.17504/protocols.io.h4qb8vw.

#### 2.2.3 LC/MS

LC/MS was undertaken by the Medical Research Support Center, Graduate School of Medicine, Kyoto University with a quadrupole–time-of-flight (Q-Tof) mass spectrometer TripleTOF 5600 (AB Sciex Pte., Ltd., Concord, Canada). Standard protocols were followed. The loading amount for each sample was 1 *μ*g. We extracted the quantitative data for the unused information for identified proteins using ProteinPilot 4.5.0.0 software (AB Sciex Pte., Ltd., Concord, Canada). For further details see [5].

## 3 Results

### 3.1 General guidelines for topological evaluations

We start from a 1-dimensional *C*^∞^ manifold with a topology, 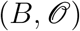. Note that many aspects of 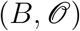 can be explained by the inverse square law by drawing on forces in the models below.

This partial topology of 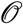 means, for example, a regular automorphism on Δ, 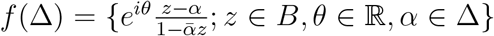 can explain anything emanating from the set of *f*, for example, isomorphism to ℝ^3^ space as shown in [1], and explored in more detail below. An apparently neutral particle system introduced with hierarchies by Galois extension could be 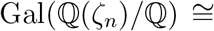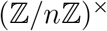 when ζ_*n*_ is a cyclotomic field. If GCD(*n*, *m*) is 1, 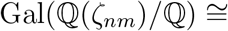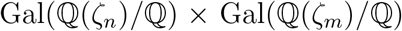. This would lead to a Kummer extension decomposed to species with *p* identity [1].

For a topology of 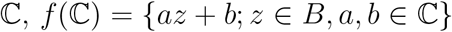 and isomorphic to ℝ^4^, later indicated as (3 + 1) dimensions with a time dimension. Obviously interaction of a complex metric, e.g. *s*^2^, *w*^2^ in [1], can induce a time dimension. For a topology of 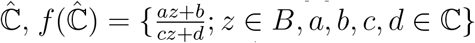 and isomorphic to ℝ^6^(ℝ^3^ × ℝ^3^), later indicated by letting ℝ^4^ compact by inducing a hierarchy as in [1].

Fundamentally, a simply connected subregion without holes such as a Riemann surface induced during hierarchization is isomorphic and holomorphic to either Δ, ℂ, or 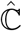. Schwarz-Christoffel mapping enables a conformal transformation from polygons to one of those regions, and the Widely Applicable Information Criterion (WAIC) has a central role as an analogy to logarithmic velocity in fluid mechanics calculated from *D* [1]. Without singularity, this is straightforward to consider and we focus on the case for singular points. As in the Bethe ansatz [2], a single dimension *z* with a particular topology is able to induce both a (3 + 1)-dimensional system and hierarchies.

### 3.2 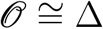 case

The Riemann-Roch theorem states

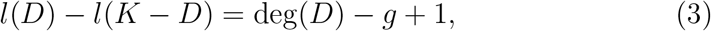

where *D* is a divisor, *K* is a canonical divisor, and *g* is a genus number. Let *TB* be a bundle. An interaction, 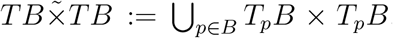, becomes a 3-dimensional *C*^∞^ manifold. Let open base elements of the manifold be *x*, *y*, *z*, and the planes on the bases be *X*, *Y*, *Z*. If we consider interactions of these bases, the left term of Eq. (3) is 3, from *g* = 10 and deg(*D*) = 12.

Let *F*

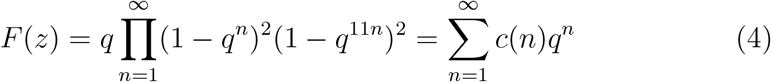

be a totally real number field of degree *g* over ℚ, and 𝕂 be a totally imaginary quartic extension of *F*. Let *D* and *D*^*int*^ be simple algebras over 𝕂 with *D* = *e*^*s/b*^. Let **G** = **GU**(*D*, *α*) with *α* being a second kind involution of *D*. Take a 3-dimensional *ℓ*-adic system in which 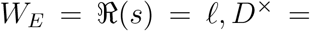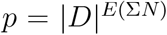, *GL*_*d*_(*E*) = *v* = ln *N_k_*/ln *p*, where *W_E_* denotes the Weil group of center *E* as a Langlands correspondence [6] [1] [5]. *ℓ* is obviously an étale (crystalline) topology independent of moduli *N_k_*, in the sense that a homomorphism of Noetherian local rings is unramified and flat, and the object is a localization of a finitely generated algebra of the origin [1]. These *p*(*ℓ*)-adic geometries are analogical to real differentiables and Clifford-Klein geometries as calculated later. The 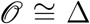 case visualizes both persistence homology *p* and étale cohomology *l*.

### 3.3 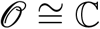 case

A Minkowski metric small *s* [1] can be utilized for a time developing model when sin, cos of the metric are converted to sinh, cosh. However, for more detailed analysis, another Minkowski metric in our model could be

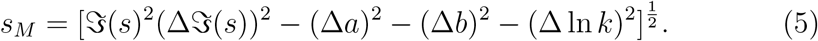

In this sense, the world line of a species is identical and a different species is non-zero, discretely depending on 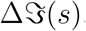. When we take 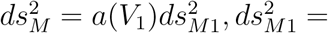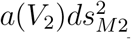, and so on. 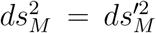 due to a Lorentz transformation and 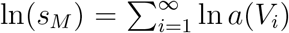 becomes a module when 2*ds*_*M*_ = 0. A set of species can thus be characterized by this module of *s_M_*. A Lagrangian could be

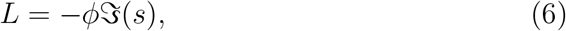

and a Hamiltonian could be

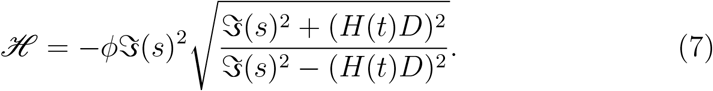

We can consider *D*′ ≅ *D_int_*, **G**′ ≅ **G^int^**, and a time dimension is induced by some admissible isomorphisms (Proposition 2.5.6 in [7]). Note that ‘temperature’ *b* and root of time *t* are closely correlated by *t* = *b* arg *D* [1]. Now consider the Poincaré conjecture, where every simply connected closed *n*-dimensional manifold *W_E_* is homeomorphic to *n*-dimensional sphere *S^n^*.

Let a Morse function be *f*: *W_E_* → [*a*, *b*], in which *a*, *b* are regular values. Let *f* have critical points *p*, *p*′ that correspond to indexes λ, λ + 1 as time. Consider that *S*^*n*−λ−1^ and *S*^λ^ cross at a single point; this indicates the status of present. The exchange of Morse functions would result in no new critical point appearing and disappearance of critical points *p*, *p*′ (*h*-cobordism theorem). This is what happens at the present state following the time arrow. Remark that *p*, *p*′ are linked to a Hecke ring via non-trivial zero points of Riemann zeta [1], fulfilling the condition of the Yang-Baxter equation. Thus this phenomenon is closely related to an analogy to quantum entanglement and face models [9, 8]. Of course, in the case of species, as species still exist, they will reappear with different p values in this model.

In this sense, for any labelling of time points 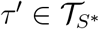, a potential for the Petersson-Weil metric is as follows:

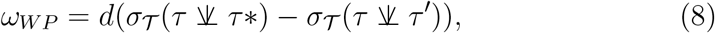

when 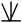 is a quasi-Fuchsian Kleinian group [10]. The ‘mating’ represents the coupling of times corresponding to *p*, *p*′.

Now consider *p*, *p*′ as characteristics on a field *k*, as in *d* = *p* = 0 in [5]. Let *E* be a singular hyperelliptic curve of the system. Real *D* will be a tensor product of an endomorphism of *E* on 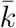 and ℚ, approximately. The resultant *D* is a quaternion field on ℚ. Take a set of ln N as an *ℓ*-adic rational Tate module as in [5]. *D* will only ramify at *p*, *p*′ or a point at infinity (c.f. [11]). This restricts the possible direction of the time arrow to vanish *p*, *p*′ only.

Generally, for species, we draw a picture of time development when the observer is at *k* = 1. For other observations, we can simply take *k* → *k*′ shifts for the calculations. That is, we can take a cyclotomic field related to the number of *k*_max_. In this sense, time in the context of a complex metric can be utilized and the world line is in web form branched at each cross-section of *p* and *p*′, not in parallel as discussed in some studies. For moving one distinct world line to another, we need velocity 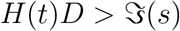.

Next, shift from *p* to 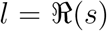 following the method outlined in [1], and simply consider a combinatory function in a probability space, Γ(*s* + 1) = *s*Γ(*s*). This is an example of a shift map. If we take a function similar to a Γ function, we can observe discrete time development merely by multiplying a master s function if we know the particular *s*. That is, adding a single fractal dimension in the past world (subtracting a single dimension from the future world by an observation) results in a simple multiplication of *s* and master Γ(*s*). Therefore, only evaluating an *s* of interest is sufficient for this aim.

Similarly, consider the Maass form of the Selberg zeta function in [1] as calculating the mode of species dynamics. Stirling’s approximation would be 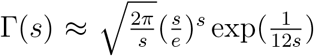, and considering a first-order approximation of the exponent with (1 + 1/12*s*) can suitably approximate the situation with superstring theory of 12 dimensions. For further approximation, we need additional dimensions. Jacobian mapping independent of a path λ

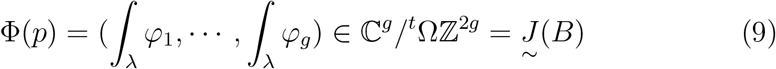

is one choice. If we know the master Maass form as the invariant form for *ρ_G_*(*c_G_*) = *c*_*G*_*Id*_*W*_ when *Id*_*W*_ is an identity mapping of a system of interest (Stone-von Neumann theorem; [12, 14, 15, 13]), differential operation does not cause any difference in the form. This ensures the condition for a suitable 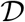-module and the accompanying derived category. Thus we can adopt a modified microlocally analytic *b* function as ∂_*b*_ = *i*∂ as a substitute for the differential operation; i.e., 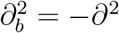, rotating the form in the angle of π, and 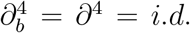, reverting back to the original orientation of the form. An Ornstein-Uhlenbeck operator would be 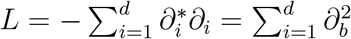. Setting a bounded Baire function *h* on ℝ^*d*^ and *f* as a solution of *Lf* = *h*− < *h* >, 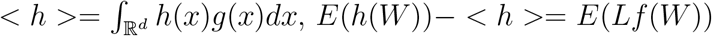 means a deviation from the expected function *h* value in the future. The operator ∂_*b*_ thus characterized for an operator calculating a future state. 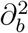 could be an element of a 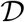-module as 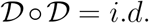 Then ∂_*b*_ would develop to analogies to energy or momentum, ∂_*b*_/∂_*t*_ = *E* or 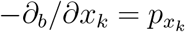 as variations of operators. The π/2 rotation of 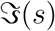 in [1] is thus justified by the modified *b* function. Considering (3 + 1) dimensions with an interaction of two 2-dimensional particles, this theory and transactional interpretation of quantum mechanics [16] are suitable. If we regard 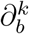, *k* ∈ ℤ as ideals of a finitely generated Jacobson radical, Nakayama’s lemma shows maintaining identity before and after the operation means the module is zero. Therefore, in this finite case, everything is an observant and at least an infinite generation is required to achieve the values out of zeros. That means, if we see something, time is infinite. Hironaka’s resolution of singularities at characteristic 0 implies such a mating of *p*, *p*′.

To resolve such a master relation, consider a form of “velocity” as *v* ∈ *TB*. Then take a 2-dimensional space consisting of *s* ∈ *B*. *s*(*v, t*) = *p*(*v*) + *tq*(*v*) as in a Lagrange equation. The Gauss curvature of this surface *K* ≤ 0. *K* ≡ 0 is only achieved when *TB* is time-independent, and this *TB* with *K* = 0 is the time-invariant bundle usable for 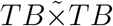 calculation for a 3-dimensional system and 6-dimensional hierarchies. Additionally, the Legendre transformation of the above equation is *X* = *v*, *Y* = *tv* − *s*, *Z* = *t* and 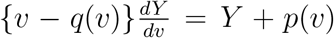. *K* = 0 means *v* = 0 and *s* = *p*(0) is the required solution. Furthermore, *s* can be regarded as a Dirac measure (*w* is a counterpart of mass and *s* = *w* + 1), and *s*′ = −*s* can be regarded as a Schwartz distribution. Although addition is allowed in the distribution, generally multiplication is not (we will illustrate that it is feasible later). However, setting the differential as 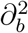, it becomes first order with a minus sign and differentiation by time: *t*^2^ is plausible. For instance,

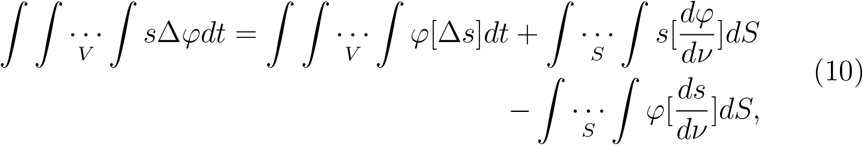

where *φ* is a distribution of interest, *s* ∈ *S*, and *ν* is a differential by unit area. The first term on the right is noise, the second is related to fractal structure, and the third is oscillative behavior. Besides singular points, it is regular. An entire function considering negative even singular points of *l* − *n* regarding *w* = *s* − 1 would be

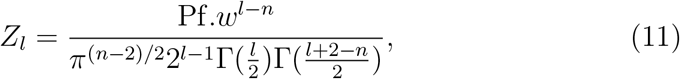

where at the singular points, *k* ∈ ℤ_≥0_, *Z*_−2*k*_ = ◻^*k*^*w*; 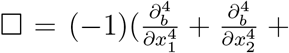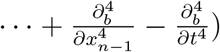. In the ∅ =∂*B* case, ◻*Z*_2_ = *w*, ◻^*k*^*Z*_2*k*_ = *w*. This means, periodical population bursting/collapsing by negative even *w* values [1]. For negative odd *w* values, chaos ensues (Šarkovski, Stefan, Block theorem) [17]. Thus, adopting *s*, *w* is suitable for applying a single-dimensional model. *s* is a measure provided it is finite in bounded domains. Therefore, singular points reflect appearance/disappearance of fractal structures. In summary, a topology 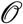 should be ({*m* = *k*} ⊂ ℕ, {ε = *b*}, {Ω = *a*}) of *N_k_* = *a* − *b* ln *k* in [1]. For further details regarding distributions, see [18].

Now let *E* be an elliptic curve: *y*^2^ + *y* = *x*^3^ − *x*^2^ as in [19]. This is equivalent to *y*(*y* + 1) = *x*^2^(*x* − 1). If we consider (3 + 1)-dimensional *N* = 1 *SU* (2) without fluctuation, *x*^2^ could be mass, (*x* − 1) could be a goldstino as spontaneous breaking of supersymmetry, *y* could be 3-dimensional fitness *D* with fluctuation, and *y* + 1 could be (3 + 1)-dimensional *s* [1]. The goldstino would represent temporal asymmetry. In Gaussian ensembles, a complex system GUE breaks time-reversal and a self-dual quaternion system GSE preserves it. Therefore *y* + 1 preserves time symmetry and consequently the present *y* breaks the symmetry.

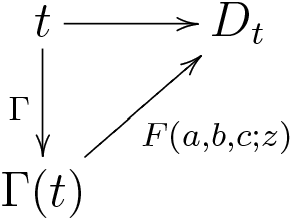

A Riemann scheme would uniformize the fitness space as a hypergeometric differential equation.

Now consider

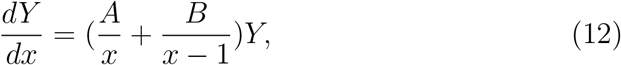

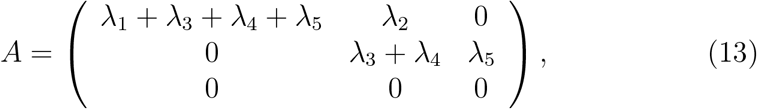

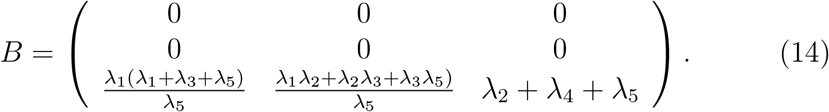

This will culminate in a generalized hypergeometric function _3_*F*_2_ that satisfies a Fuchs-type differential equation _3_*E*_2_. If we set proper region Δ (13 different regions),

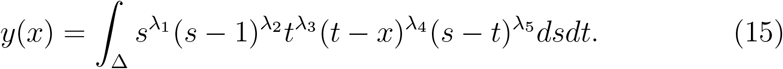

*x* = 0, *w* = *D*, *s* = 1 would result in

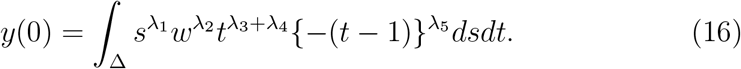

λ_1_ = λ_2_ = λ_3_ = λ_4_ = λ_5_ = 1 would be *E*^2^: − ∫*y*(*y* + 1)*x*^2^(*x* − 1)*dxdy* form, obviously the integral of the interaction of two elliptic curves.

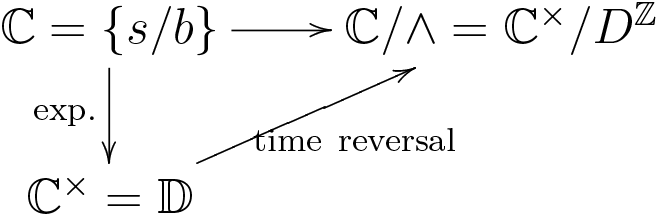

For consideration of an interacting 4-dimensional system, let us take Painlevé VI equations on a (3 + 1)-dimensional basis with a single Hamiltonian [20] [21]. The Hamiltonian should be 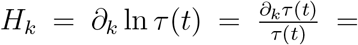 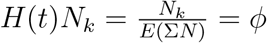 when *H*(*t*) is a Hubble parameter [22] [1]. *τ* (*t*) is thus an inverse of a Hubble parameter, and its kth boundary is a *k*th species. Note that the 3-dimensional system represents the smallest possible number of dimensions whose associativity equations become non-empty even in the presence of the flat identity. Furthermore, considering a fundamental group π_1_ of *C*_0,*n*_:= ℙ^1^\{*z*_1_,…, *z_n_*}, the dimension of representations *ρ* of π_1_ in *SL*(2, ℂ) is 2(*n* − 3) [22]. If we would like to set π_1_ as an étale topology with 0 dimension, *n* = 3. (3+1)-dimensional semisimple Frobenius manifolds constitute a subfamily of Painlevé VI:

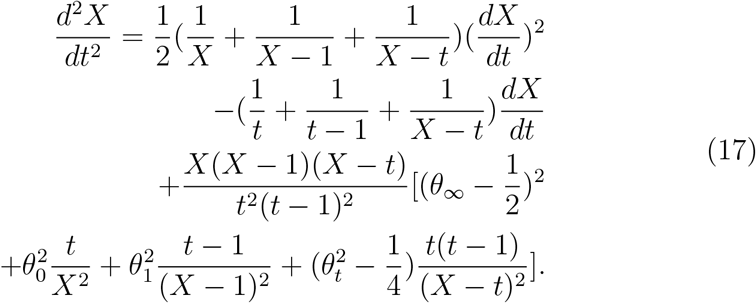

Recall that the above equation is related to a rank 2 system:

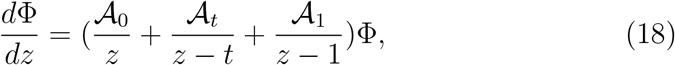

or

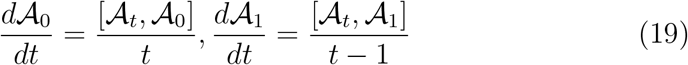

with 4 regular singular points 0, *t*, 1, ∞ on ℙ_1_. Also,

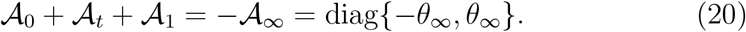

Note that the total sum of the matrix system is equal to 0. Assuming a 3-wave resonant system [23],

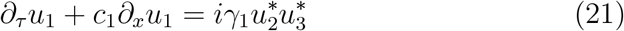

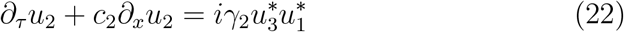

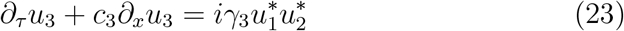

An expansion of this model results in the 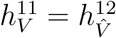 mirror symmetry relation for the Calabi-Yau threefolds. Recall that matrix Painlevé systems of two interacting systems

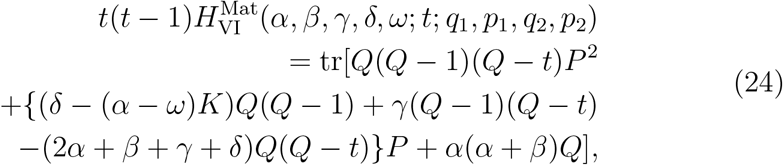

has 11 parameters.

Now let us convert a Painlevé VI equation to a more realizable form as in physics. The Painlevé VI equation is equivalent to

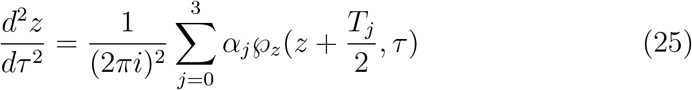

where 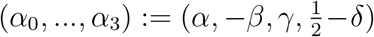, (*T*_0_,…, *T*_3_) = (0, 1, *τ*, 1+*τ*), and *℘* is the Weierstraß*℘* function (Theorem 5.4.1 of [20]). Furthermore, any potential of the 3-dimensional normalized analytic form

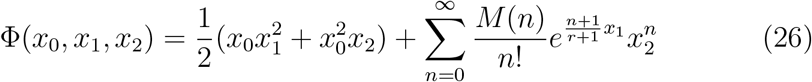

can be expressed through a solution to the Painlevé VI equivalent with 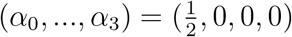, that is,

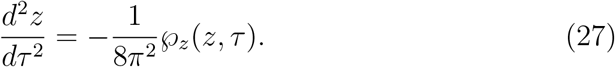

When *q* = *D* = *e*^*iπτ*^, the Picard solution of the *τ* function on the 4 dimensions that corresponds to the *c* = 1 conformal field blocks in an Ashkin-Teller critical model would be

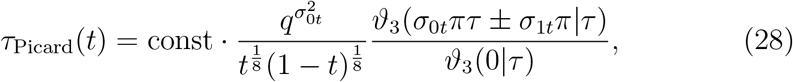

where the Jacobi theta function is 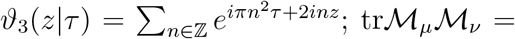 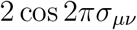 when the parameter space of (*θ*_0_, *θ*_*t*_, *θ*_1_, *θ*_∞_) is 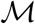 [24] [25] [22] [26]. For other algebraic solutions, see [27]. Let us calculate a Clifford algebra in an *n* = 3 system [28]. First, let the representation (*ρ*, *V*) of the algebra *Cl_n_* fulfill the condition *ρ*: *Cl_n_* ∍ *ϕ* ⟼ *ρ*(*ϕ*) End(*V*) with *ρ*(*ϕ*)*ρ*(*ψ*) = *ρ*(*ϕψ*). When *n* is odd, for example, 3, there are nonequivalent representations:

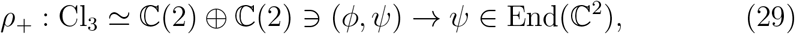

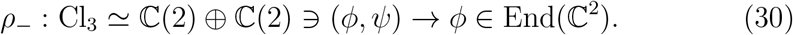

For example, let us calculate a complex *v*, *v*′ by 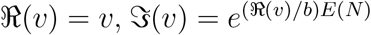, 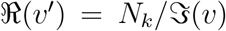, and 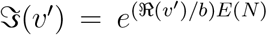 as in [5]. The next complex *v*′′ is 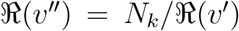 and 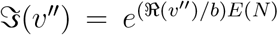. We can calculate *v*′′′ by the same operator as before. We denote this situation RRR. Graphing the calculated 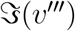 values with their rank among 800 proteins permits classification into 3 groups demarcated based on slope values, namely, values below 1.01, between 1.01 to 2.00, and above 2.00 (Fig. 1). The 0.30 value of Filamin-A was excluded because it probably mostly reflects adapted proteins in fibroblasts (HEK-293). The irreducible representations in the raw LC/MS data of [5] are 4-dimensional 1–2 (average 1.368 ± 0.004, 99% confidence) in non-adapted situations and 3-dimensional 1 (average 1.001571 ± 0.000006, 99% confidence) in adapted situations, respectively (Supplementary Table 1). The remainder are probably repressed (disadapted) proteins. In tensor algebra 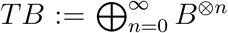, *B* = ⨁_*i*∈*I*_ *RX*_*i*_, *x* ∈ *X*, *x* ⨂ *x* −*q*(*x*) ∈ *R* ⨁ *B*^⨂2^, *x* is a single fractal dimension (= *w*), and the fractal dimension of *q*(*x*) is 1/2, 1 for non-adapted and adapted stages, respectively [1]. We are thus able to calculate a characteristic number related to protein adaptation.

**Figure 1:**
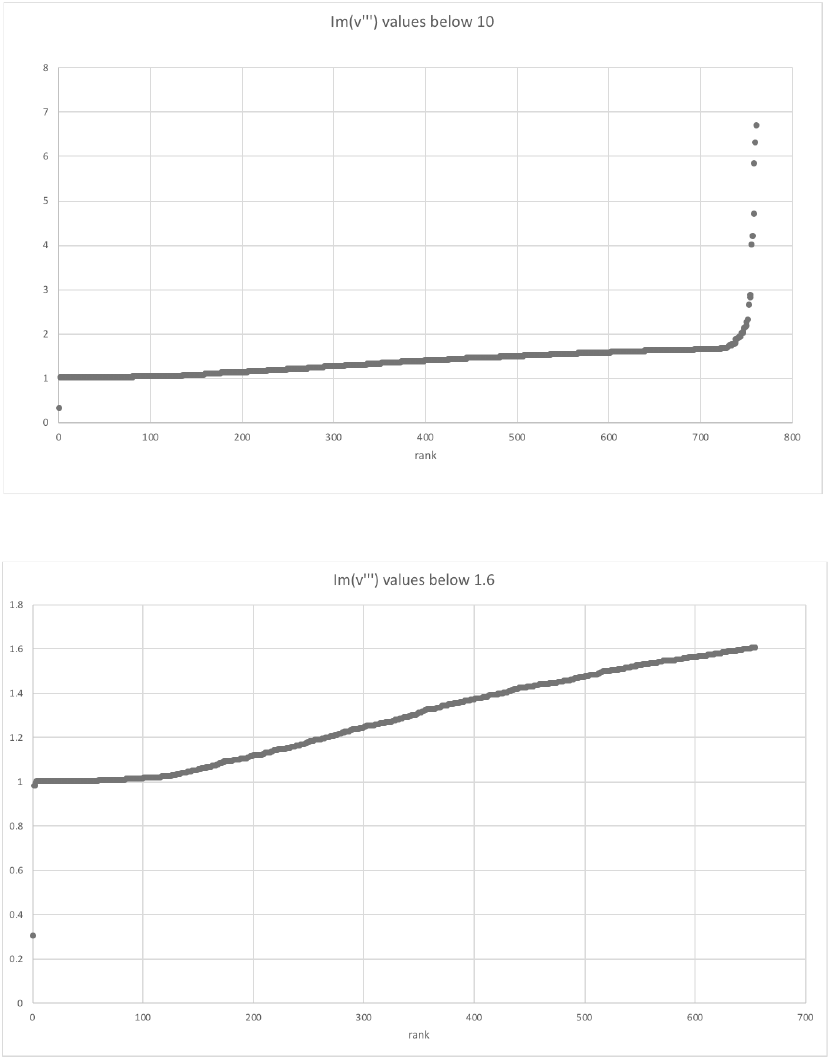
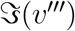 values versus their ranks.

### 3.4 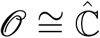 case

For the species data set (Table 1) [1], consider that a sequential operation is an exact form. As in [5], setting operation III as 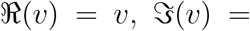 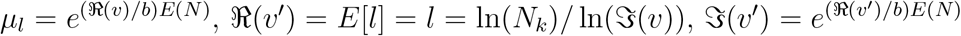 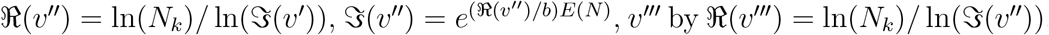 and 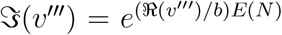, we have 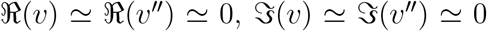, 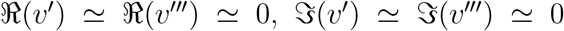 (Table 1), suggesting that an actual/potential of species creates an actual/potential appearance of the adapted hierarchy above two layers. Recall that this is a short exact sequence; the morphism 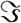 becomes monomorphism and 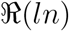 becomes epimorphism. Furthermore, 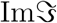 is equal to 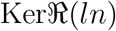. Obviously there also exists a homomorphism 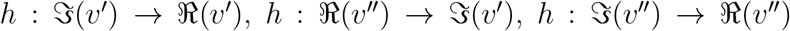 or 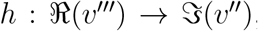, and the short exact sequence is a split. These are abelian groups and 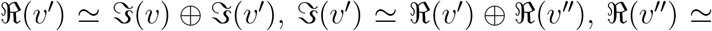 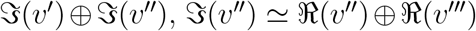. The data show that an actual layer is a direct sum of a potential layer below and a potential layer. The data also show that a potential of the layer is a direct sum of a real layer and a layer above the layer. Finally, defining a Galois action Gal(*L/K*), actions defined by 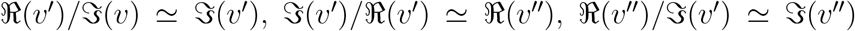, and 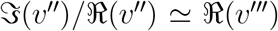 are all Galois, achieving our goal for defining proper Galois actions with a topology of *v* for biological hierarchies. A species is thus likely to emerge from the interaction of species.

**Table 1:** *N* values.

| $N$ | $P. pallidum$ (WE) | $D. purpureum$ (WE) | $P. violaceum$ (WE) |
| --- | --- | --- | --- |
| May | 0 | 76 | 0 |
| June | 123 | 209 | 52 |
| July | 1282 | 0 | 0 |
| August | 1561 | 0 | 0 |
| September | 901 | 107 | 0 |
| October | 1069 | 35 | 0 |
| November | 60 | 0 | 101 |
| December | 190 | 0 | 0 |
| January | 29 | 0 | 0 |

| $N$ | $P. pallidum$ (WW) | $D. purpureum$ (WW) | $P. violaceum$ (WW) |
| --- | --- | --- | --- |
| May | 0 | 83 | 0 |
| June | 147 | 0 | 0 |
| July | 80 | 215 | 320 |
| August | 1330 | 181 | 0 |
| September | 809 | 77 | 649 |
| October | 799 | 0 | 107 |
| November | 336 | 0 | 0 |
| December | 711 | 0 | 0 |
| January | 99 | 0 | 0 |

WE: Washidu East quadrat; WW: Washidu West quadrat (please see [3]). Scientific names of Dictyostelia species: $P. pallidum$ : *Polysphondylium pallidum*; $D. purpureum$ : *Dictyostelium purpureum*; and $P. violaceum$ : *Polysphondylium violaceum*. $N$ is number of cells per 1 g of soil. Species names for Dictyostelia represent the corresponding values. Red indicates $\Re(s)$ values of species that were approximately integral numbers greater than or equal to 2.

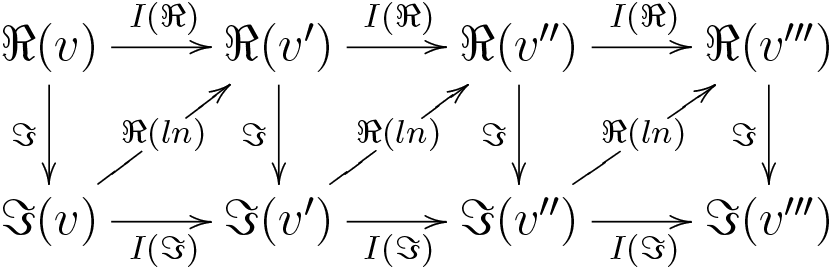

For species [1], consider that a sequential operation in the previous sections is an exact form. As in [5], setting an operation III, we have 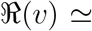 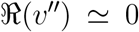 and 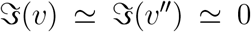, but no further (Table 1), suggesting that an actual/potential of species creates an actual/potential appearance of the adapted hierarchy above two layers, which diminishes in the three layers above. This might reflect effects from different time scales among different layers [3]. Similar to the previous section, 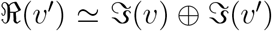 and 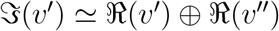.

From the III morphisms, we can draw a short exact sequence corresponding to 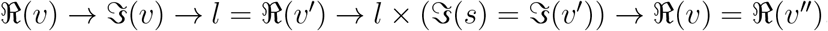,

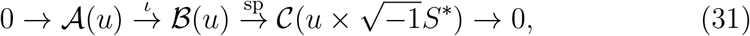

regarding *g* = *l* as a specific spectrum of the Schwartz distribution (or Sato hyperfunction [29, 30]) of a microfunction sp *g* [32, 31]. Not only addition, but also multiplication is feasible for −*s* in this regard.

### 3.5 Congruent zeta function

Hereafter we will adhere to the situation where 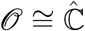. For the other aspect, instead of 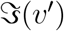, we can consider ℤ/*l*ℤ, by 1/*l*-powered 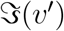, state a *p*-adic number correspondence, and then take a valuation of it. Universal coefficient theorems [33],

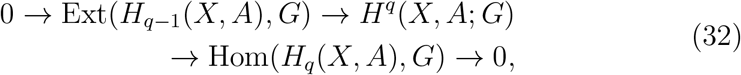

could be described as

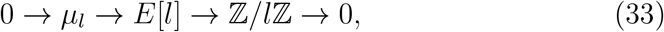

making an exact sequence, with 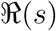 value in the middle level between populational 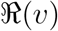 value and its fractal 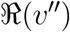 value. *E*[*l*] → ℤ/*l*ℤ is an injection and ℤ/*l*ℤ → 0 an epimorphism. The image of the former is the kernel of the latter. Homology backwards is a homomorphism of the cohomology, and the exact sequence splits. These are abelian groups and *E*[*l*] ≅ *μ_l_* ⊕ ℤ/*l*ℤ; ℤ/*l*ℤ ≅ *E*[*l*] ⊕ 0. A real level is constituted by a direct sum of a potential level below and its own potential. A potential level is constituted by a direct sum of a real level below and a real level above. *E*[*l*]/*μ_l_* = ℤ/*l*ℤ; ℤ/*l*ℤ/*E*[*l*] ≅ 0 are Galois actions and a representation of an étale topology *ℓ* is obtained, concomitantly with information of interactions among different levels of hierarchies. Species should appear two layers above the population layer. [3] reports results where the point mutation rate is on the order of 10^−8^ and speciation is on the order of 10^−25^, roughly above a square of 10^−8^ over 10^−8^. This calculation could be modeled by a simple critical phenomenon of dendrogram percolation. In this model, approaching 1/2 − 0 probability of mutation maintenance leads to divergence in cluster size. Regarding non-trivial ζ(*w*) = 0 as a seed for speciation, a ~10^8^ population is on the same order as a branch for being identical to ancestors or different from them at each genome base pair. A dendrogram can be regarded as a phylogenetic tree for dividing cells, which is common to both asexually propagating organisms and a constituent of sexually reproducing organisms at the level of cell division of germ line cells, strictly correlated to mutation during cell cycle processes. These facts exhibit *ℓ* and Galois actions can adequately describe interhierarchical interactions.

The logic above would suggest application of Grothendieck groups. Let the situation be a Noetherian ring, i.e., *B* is the ring. Let *F*(*B*) be the set of all isomorphisms of *B*-modules. Let *C_B_* be the free abelian group generated by *F*(*B*). The short exact sequence above is associated with (*μ_l_*) − (*E*[*l*]) +(ℤ/*l*ℤ) of *C_B_* (() is an isomorphism). Let *D_B_* be the subgroup of *C_B_*. The quotient group *C_B_*/*D_B_* is a Grothendieck group of *B* related to potential of *s*, *w* layers, denoted by *K*(*B*). If *E*[*l*] is a finitely generated B-module, γ(*E*[*l*]) would be the image of (*E*[*l*]) in *K*(*B*). There exists a unique homomorphism λ0: *K*(*B*) → *G* such that λ(*E*[*l*]) = λ0(γ(*E*[*l*])) for all *E*[*l*] when *G* is an abelian group of the B-module. This representation corresponds to the Stone-von Neumann theorem in this restricted situation. *B* is generated by γ(*B/p*) when *p* corresponds to species in a biological sense. If *B* is a principal ideal domain constituting a single niche without cooperation of distinguished niches, *K*(*B*) ≅ ℤ, and this is suitable when considering biological numbers for individuals. Considering different *E*[*l*], *M_l_*, and *N_l_*, and the set of all isomorphisms of a flat *B*-module *F*_1_(*B*), γ_1_(*M_l_*) · γ_1_(*N_l_*) = γ_1_(*M_l_* ⊗ *N_l_*); γ_1_(*M_l_*) · γ(*N_l_*) = γ(*M_l_* ⊗ *N_l_*); *K*_1_(*A*) ≅ ℤ with tensor products. Furthermore, if *B* is regular, *K*_1_(*B*) → *K*(*B*) is an isomorphism. The sum of interactions for different niches (not interacting between distinguished niches) is thus calculable as integers by a Grothendieck group. If the calculation does not lead to integers, the situation involves interactions among distinguished niches. Algebraic expansion of this ring thus introduces entirely different niches to the original ring. If *a* ∈ *K*, *f*(*x*) = *x^l^* − *x* − *a*, *α* ∈ *K̄*, *f*(*α*) = 0, *α* ∉ *K*(*α* ∈ ∂*K*), *f*(*x*) is irreducible on *K*, *L* = *K*(*α*) is a Galois extension, and Gal(*L*/*K*) ≅ ℤ/*l*ℤ. *α* is from the hierarchy above based on a new ideal.

To unify the sections introducing Galois *H^i^* and the preceding sections regarding the time arrow, consider *X*, *Y*, which are eigen and smooth connected algebraic curves on an algebraic closed field.

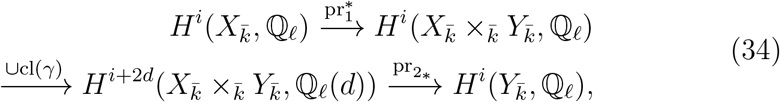

when γ is an algebraic correspondence from *Y* to *X*. If we assume *X* and *Y* correspond to different time points, the above diagram,

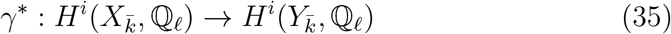

describes the time development of the system. To dissect the contributions of each component on the time developing system, let *κ_m_* be an *m*-dimensional expansion of *κ*, which is a finite field of a residue field of an integer ring *O_K_* on *K*. When the eigen smooth scheme *Y* is on *κ*,

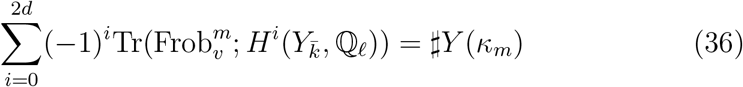

[34] [35].

When *Y* is finite, a congruent zeta function is

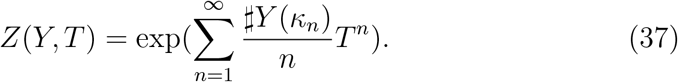

Setting

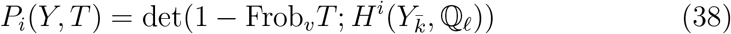

results in

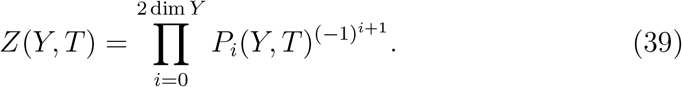

To separate each contribution of *H^i^*, consider Weil conjectures [36] [37], and *P_i_*(*Y, T*) and *P_j_*(*Y, T*) are disjoint when *i* ≠ *j*. *P_i_*(*Y, T*) and 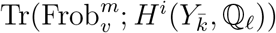 are thus calculable and this deciphers each contribution of *P_i_*(*Y, T*)s. Examples of the calculation are provided in Tables 2 & 3. Generally, large positive zeta values represent highly adapted situations, whereas large negative zeta values represent highly disadapted situations and zero values are neutral situations. *P*_0_, *P*_1_, *P*_2_ correspond to 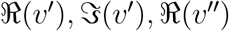, values close to zero represent large contributions, and for 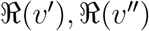, large values represent large contributions. The inverses of 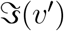 scale for 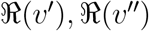. The important point here is that by utilizing a congruent zeta function, we can visualize a contribution from each hierarchy.

**Table 2:** Calculations for Washidu East quadrat

| Z (congruent) | *P. pallidum* | *D. purpureum* | *P. violaceum* |
| --- | --- | --- | --- |
| May | - | - | - |
| June | 0.009378 | 151.1 | 9.272 |
| July | - | - | - |
| August | - | - | - |
| September | 114.7 | 30.89 | - |
| October | 334.6 | -540.4 | - |
| November | 0.02561 | - | -54.13 |
| December | - | - | - |
| January | - | - | - |

| $P_0$ | *P. pallidum* | *D. purpureum* | *P. violaceum* |
| --- | --- | --- | --- |
| May | - | - | - |
| June | -1.288 | -0.06806 | -0.1520 |
| July | - | - | - |
| August | - | - | - |
| September | -1.163 | -0.7248 | - |
| October | -0.8250 | 0.02954 | - |
| November | -1.002 | - | 0.1790 |
| December | - | - | - |
| January | - | - | - |

| $P_1$ | $P. pallidum$ | $D. purpureum$ | $P. violaceum$ |
| --- | --- | --- | --- |
| May | - | - | - |
| June | 0.7635 | -10.29 | -1.253 |
| July | - | - | - |
| August | - | - | - |
| September | -133.4 | -22.39 | - |
| October | -276.1 | -15.96 | - |
| November | 0.7480 | - | -9.689 |
| December | - | - | - |
| January | - | - | - |
| $P_2$ | $P. pallidum$ | $D. purpureum$ | $P. violaceum$ |
| May | - | - | - |
| June | -63.23 | 1.000 | 0.8886 |
| July | - | - | - |
| August | - | - | - |
| September | 1.000 | 1.000 | - |
| October | 1.000 | 0.9999 | - |
| November | -29.13 | - | 1.000 |
| December | - | - | - |
| January | - | - | - |

**Table 3:**
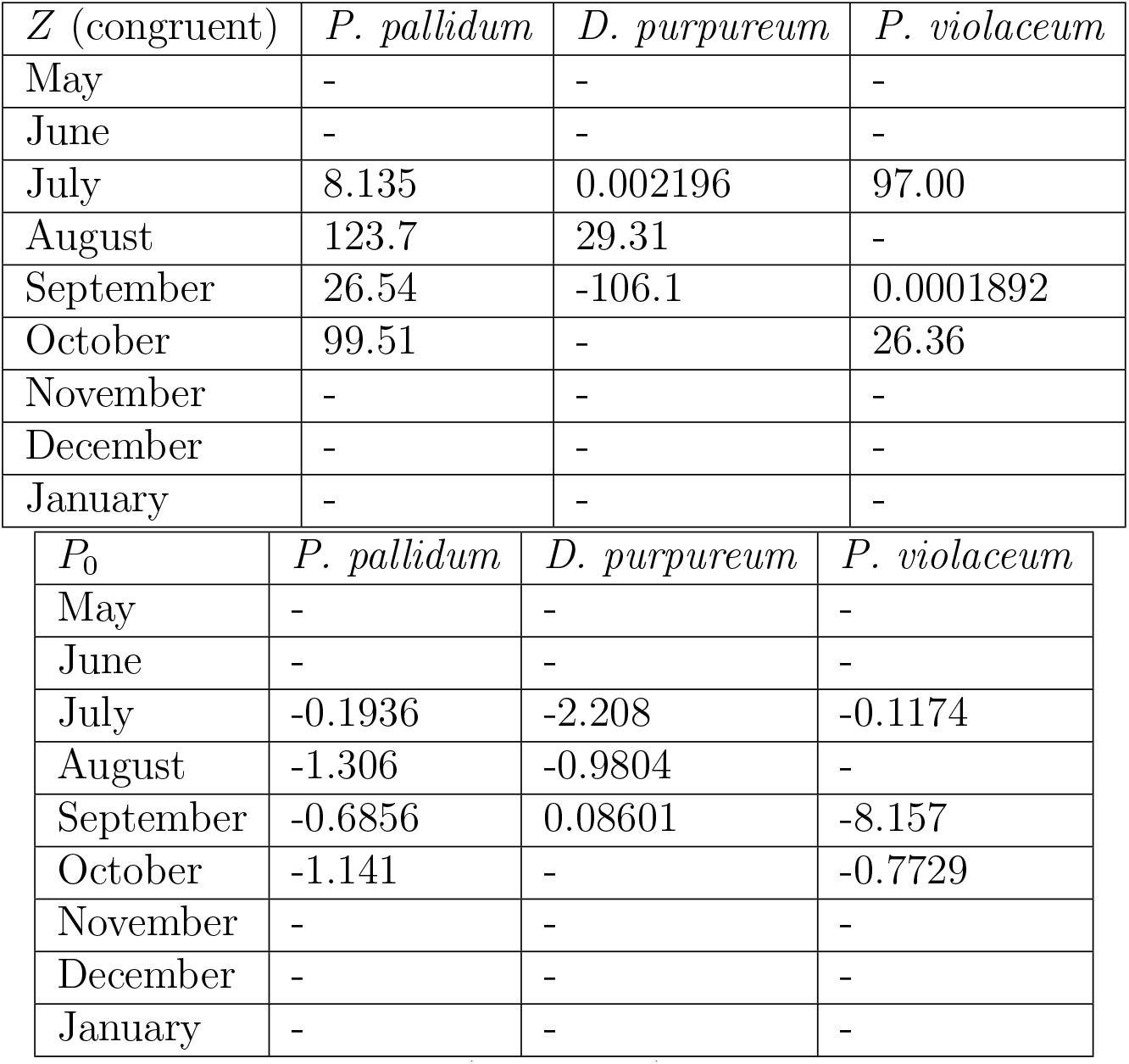
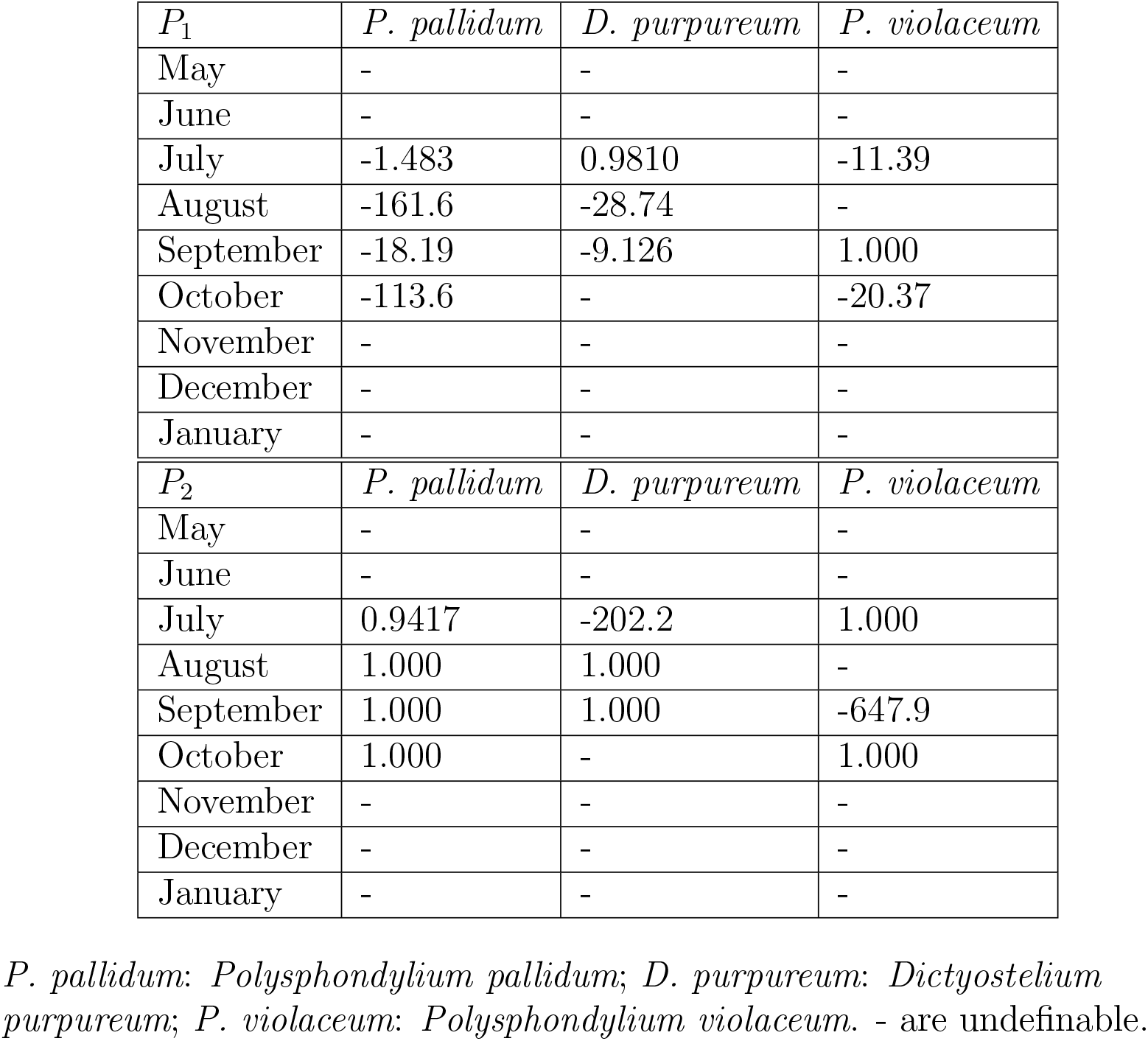
Calculations for Washidu West quadrat.

From these theorems, we can deduce that **P**^2^ is a pencil on elliptic curves with a section of order two and an additional multisection. Setting ζ = *e*^2π*i*/3^ = (*e*^π*i*/3^)^2^ on the initial condition of **P**^2^ at the point *x_a_* = 0,

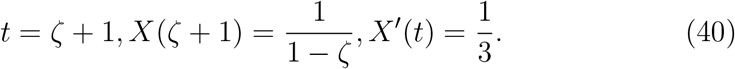

In the PzDom model [1], 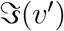 for predicting the future and *t* is an addition of 1 to interactive (*e*^π*i*/3^)^2^ if 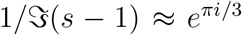 is neglectable. When in close proximity to trivial zero points of Riemann ζ, *t* ~ 1 and *X*(*t*) ~ 1. 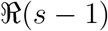 thus represents a (2 + 1)-dimensional system.

System dimensions are thus reduced to 2+1. For reproducing the kernels, let *q* be in (*Q*^∞^)^Γ^(**H**\*). Then,

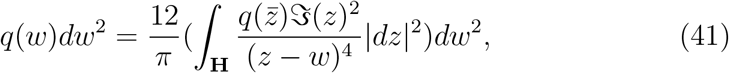

where *w* = *α*/*β* and 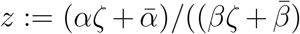. The term in parentheses is the reproduced kernel (Prop. 5.4.9 of [10]).

Now consider *q* difference Painlevé VI with 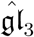 hierarchy. *q* could be equal to −*s*, and 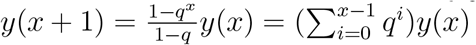 can be converted from *q* to −*s*, when *x* → ∞

Setting |*q*| > 1, *t* as an independent variable, and *f*, *g* as dependent variables,

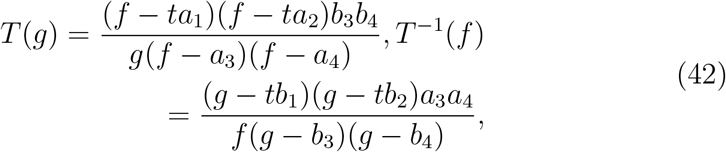

where

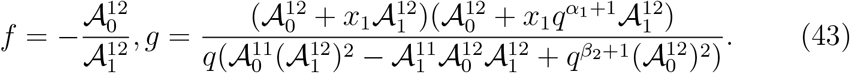

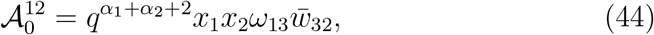

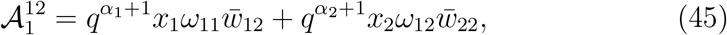

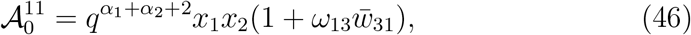

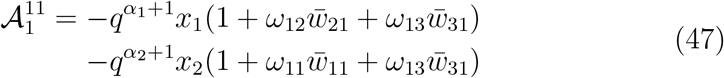

and considering *q* = −*b* ln *D* of the PzDom model [1], local time development can be easily calculated. (*a*_1_, *a*_2_, *a*_3_, *a*_4_); (*b*_1_, *b*_2_, *b*_3_, *b*_4_) have 4 parameters interacting with each other in this soliton equation of similarity reduction [38] [23]. In other words, we are treating a direct sum of two Virasoro algebras, or a Majorana fermion and a super-Virasoro algebra [25].

### 3.6 Further consideration of 1+1 dynamics

There is another way of considering system dynamics with *q*, starting from a Young tableau. Let S be a finite or countable set, for example, as the measures of species density as Specℤ. For 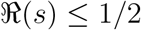, let an absolute value of an absolute zeta function 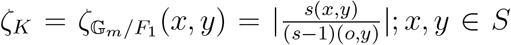 where 𝔾_*m*_ = *GL*(1). For 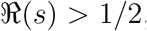, and let an absolute value of an inverse of an absolute zeta function 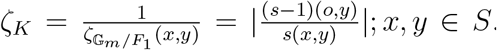. ζ_*K*_ becomes a Martin kernel. Let a distance function *D*_δ_(*x, y*) = ∑_*z*∈*S*_ *C*_*z*_(|ζ_*K*_(*z, x*) − ζ_*K*_(*z, y*)| + |δ_*zx*_ − δ_*zy*_|), where δ is the delta function. For a distance space (*S*, *D*_δ_), a topology of *S* determined by *D_δ_* is a discrete topology and (*S*, *D_δ_*) is totally bounded. A completion of (*S*, *D_δ_*) will be set as 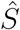. Let a Martin boundary 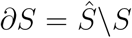 be a (*d* − 1)-dimensional species density not restricted to a random walk or transition probability. *S^d^* represents all possibilities of *S*^*d*−1^ with a time dimension. Furthermore, a set of *S*^*d*−1^ can be expressed by a Young tableau in a Frobenius coordinate system. Taking a Maya diagram of the tableau distributes the data to a single dimension. Therefore, the 3-dimensional system is in fact represented as a 1-dimensional system, a set of *F*_1_ = *F_q_*. In this context, a set of the individual numbers of species is over ℤ and a time *X* is a flat algebra Λ-space over ℤ. A Λ-structure on *X* is *ψ_p_*: *X* → *X*, where *ψ* is *X* ×_Specℤ_ Spec𝔽_pc_. In other words, Λ = ℤ[Gal(ℤ/𝔽_*pc*_)]. *p_c_* = 1 when there is no hierarchy/period in our analysis and, for example, *p_c_* = 2 in protein or species data sets described above. Therefore, the hierarchy extends from *F*_1_ to *F*_2_. 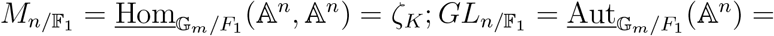 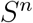 and thus *s* ∈ 𝔾_*m*_ and *s* − 1 ∈ *F*_1_ when 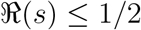 and *s* − 1 ∈ 𝔾_*m*_ and *s* ∈ *F*_1_ when 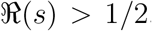. *q* ∈ 𝔾_*m*_ and Spec(*q*) is Spec(*s*) or Spec(*s* − 1). Since *D* = *e*^*s/b*^ is calculable in [1] with a root of time *t*, temperature *b_t_* at time point *t*^2^ and temperature *b*_*t*−1_ at time point (*t* − 1)^2^ when time is properly scaled, the dynamics of *q* can be calculated by this basal information. See [39] for further details in this respect as relates to Grothendieck’s Riemann Roch theorem. This is another explanation as to why a 1-dimensional system with a certain topology leads to 3 + 1 dynamics.

### 3.7 *℘* as evaluations for interactions

Take Wallis’ formula:

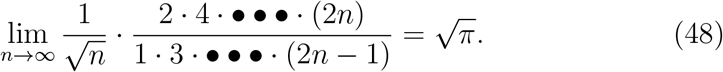

The upper product of even numbers could be a product of bosonic multiplications, and the lower product of odd numbers could be that of fermionic multiplications. The square of them divided by n as an average number of actions would result in π. π is thus the number ratio of boson multiplications and fermion multiplications. In other words, an area of a circle corresponds to boson actions and the square of the radius corresponds to fermion actions. Globally there are ~ 3 times more bosonic actions than fermionic actions. For further expansion for the bosonic even −*w* (without *w* = 0) with *μ*(*n*) = 1 [1], Weierstraß 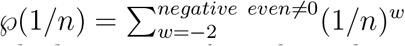 and a ((*w*/2 + 1) × *n*) (*n* × 1) matrix would calculate a set of patch quality *P_w_* of bosons involving a future status of *w* = −2. Similarly, even −*s* with *μ*(*n*) = −1 [1], 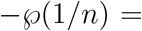 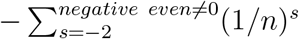, and a ((*s*/2 + 1) × *n*) (*n* × 1) matrix would calculate a set of patch quality −*P*_s_ of fermions involving a future status of *s* = −2. Regarding *w* = *s* − 1, *P*(*w*) = *P_w_* − *P*_*s=w*+1_ = ζ(*w*)+ *n* + *n*^2^ and the Riemann ζ function can be related to patch quality. Population bursts with these even *w* (odd *s*) could be calculated by *P_w_* → +∞ with negative even *w* (negative odd *s*), or in lower extent of bursting, *P_s_* → ∓∞ with *w* → 1 ∓ 0(*s* → 2 ∓ 0). Since *P* (0) ≠ 0 and *P* (0) → +∞, considering *P* (*w*) = *℘*(1/*n*) + *℘*(1/*n*)/*n* and *a_k_*, *b_k_* as zero points and poles of the function,

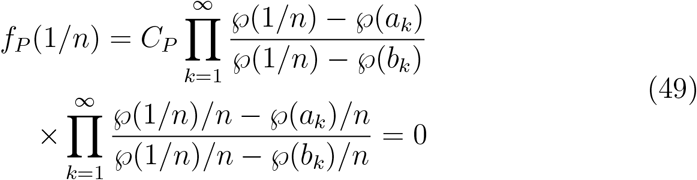

because the constant *C_P_* = 0 when *w* = 0 [40]. Thus *w* = 0(*s* = 1) means every singularity can be considered as a zero ideal adopting *f_P_*. *w* → 0 means a general limit of 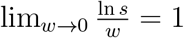. We can regard a logarithm of *s* as a fitness when the fitness is sufficiently small. A fixed point of the observer at *s* = 1 implies everything combined to the zero ideals. If we regard Weierstraß 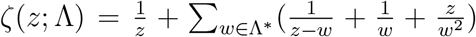 (not Riemann zeta) as a distribution *s*_1_, *s*_2_ results in

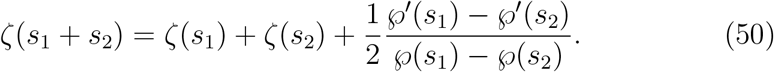

This means the third term on the right is a contribution of different fractal hierarchies, besides a direct sum of distribution functions. Tables 4 to 8 present values for the Weierstraß zeta function, Weierstraß *℘*, *℘*′, and interaction terms. Note that at Washidu West in September, Pv-Dp-Pp interacted strongly in that order. In October, there is also a strong interaction of Pv-Pp. Compared with Washidu West, Washidu East exhibited weaker interaction and was dominated by Pp. It is also notable that the strengthes of hetero-interactions were generally weaker than those of homo-interaction, as supposed in [1]. Of course, utilizing population data instead of species data elucidates similar or larger values for hetero-interaction terms compared with homo-interaction terms (data not shown), as was expected in [1].

**Table 4:** Weierstraß ζ values.

| Weierstraß $\zeta$ | WE *P. pallidum* | WE *D. purpureum* | WE *P. violaceum* |
| --- | --- | --- | --- |
| May |  |  |  |
| June | 2.290e15 - 5.081e15*I | 5.648e51 + 1.513e52*I | 3.036e32 + 1.2783e32*I |
| July |  |  |  |
| August |  |  |  |
| September | -9.284e28 - 2.716e28*I | -1.501e23 + 3.448e23*I |  |
| October | -3.307e36 - 2.666e37*I | -1.220e35 - 2.047e35*I |  |
| November | 3.579e14 - 1.003e15*I |  | 2.065e59 + 9.395e59*I |
| December |  |  |  |
| January |  |  |  |

| Weierstraß $\zeta$ | WW *P. pallidum* | WW *D. purpureum* | WW *P. violaceum* |
| --- | --- | --- | --- |
| May |  |  |  |
| June |  |  |  |
| July | 2.329e35 + 1.735e35*I | 7.052e8 - 2.352e10*I | 4.950e53 + 1.630e54*I |
| August | -4.121e28 - 1.547e28*I | -1.075e22 + 3.286e22*I |  |
| September | 1.493e39 + 1.008e39*I | 6.076e48 + 1.023e49*I | 1.220 + 0.02924*I |
| October | -4.379e28 - 1.562e28*I |  | -1.440e22 + 4.328e22*I |
| November |  |  |  |
| December |  |  |  |
| January |  |  |  |

WE: Washidu East quadrat; WW: Washidu West quadrat; *P. pallidum*: *Polysphondylium pallidum*; *D. purpureum*: *Dictyostelium purpureum*; *P. violaceum*: *Polysphondylium violaceum*. Weierstraß $\zeta$ are calculated from $\wp$ on an elliptic curve $[0, 1]$ , expanded to 30-th order. Constants of integration were neglected for $\zeta' = -\wp$ .

**Table 5:**
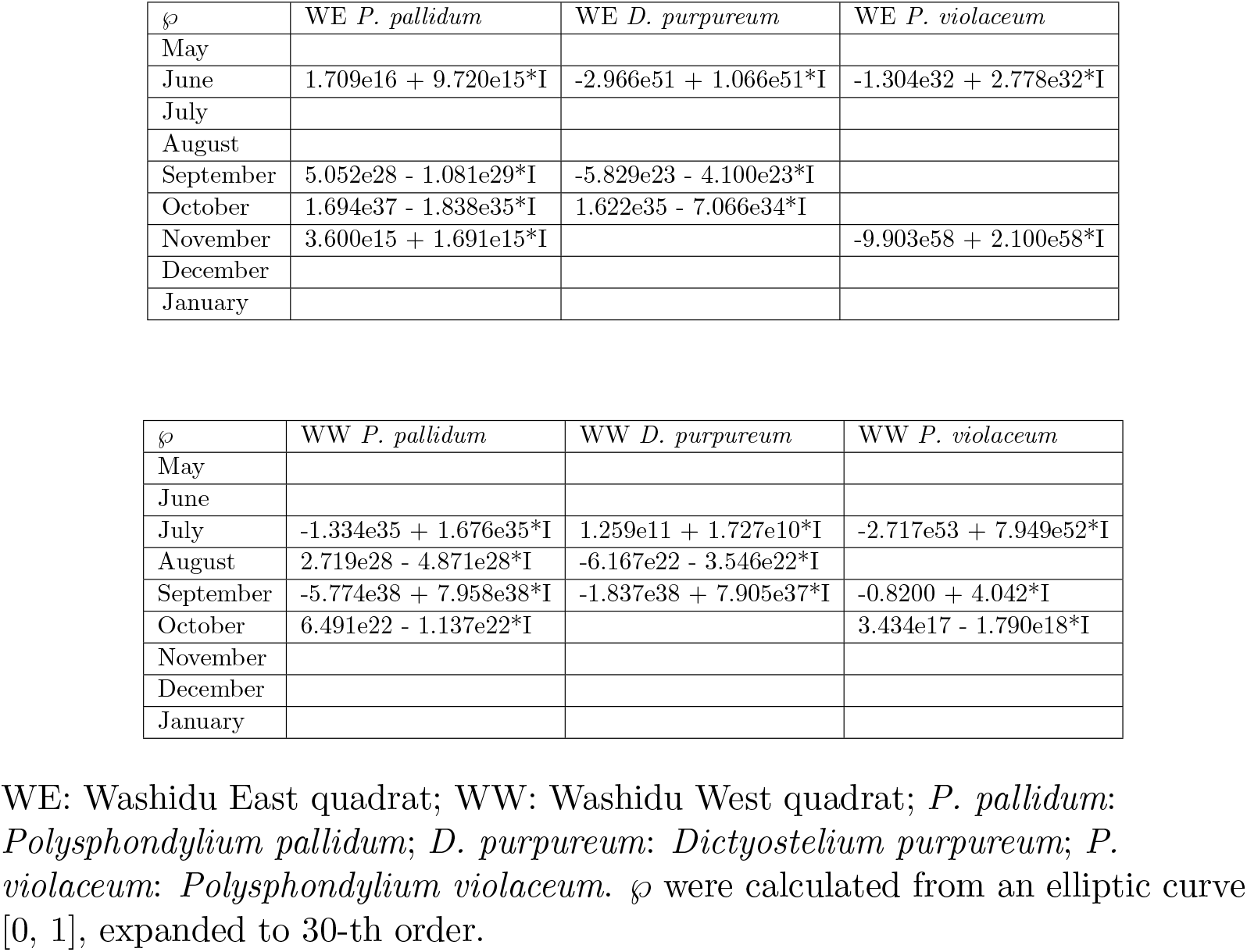
*℘* values.

| $\wp$ | WE *P. pallidum* | WE *D. purpureum* | WE *P. violaceum* |
| --- | --- | --- | --- |
| May |  |  |  |
| June | 1.709e16 + 9.720e15*I | -2.966e51 + 1.066e51*I | -1.304e32 + 2.778e32*I |
| July |  |  |  |
| August |  |  |  |
| September | 5.052e28 - 1.081e29*I | -5.829e23 - 4.100e23*I |  |
| October | 1.694e37 - 1.838e35*I | 1.622e35 - 7.066e34*I |  |
| November | 3.600e15 + 1.691e15*I |  | -9.903e58 + 2.100e58*I |
| December |  |  |  |
| January |  |  |  |

| $\wp$ | WW *P. pallidum* | WW *D. purpureum* | WW *P. violaceum* |
| --- | --- | --- | --- |
| May |  |  |  |
| June |  |  |  |
| July | -1.334e35 + 1.676e35*I | 1.259e11 + 1.727e10*I | -2.717e53 + 7.949e52*I |
| August | 2.719e28 - 4.871e28*I | -6.167e22 - 3.546e22*I |  |
| September | -5.774e38 + 7.958e38*I | -1.837e38 + 7.905e37*I | -0.8200 + 4.042*I |
| October | 6.491e22 - 1.137e22*I |  | 3.434e17 - 1.790e18*I |
| November |  |  |  |
| December |  |  |  |
| January |  |  |  |

WE: Washidu East quadrat; WW: Washidu West quadrat; *P. pallidum*: *Polysphondylium pallidum*; *D. purpureum*: *Dictyostelium purpureum*; *P. violaceum*: *Polysphondylium violaceum*. $\wp$ were calculated from an elliptic curve $[0, 1]$ , expanded to 30-th order.

**Table 6:** *℘*′ values I.

| $\wp'$ | WE *P. pallidum* | WE *D. purpureum* | WE *P. violaceum* |
| --- | --- | --- | --- |
| May |  |  |  |
| June | 3.841e16 - 5.488e16*I | 1.938e50 + 5.613e50*I | 2.450e32 + 1.2734e32*I |
| July |  |  |  |
| August |  |  |  |
| September | -1.181e29 - 7.904e28*I | -9.494e23 + 8.939e23*I |  |
| October | 1.048e36 - 1.027*I | -3.553e34 - 1.217e35*I |  |
| November | 7.322e15 - 1.234e16*I |  | 2.058e57 + 1.008e58*I |
| December |  |  |  |
| January |  |  |  |

WE: Washidu East quadrat; *P. pallidum*: *Polysphondylium pallidum*; *D. purpureum*: *Dictyostelium purpureum*; *P. violaceum*: *Polysphondylium violaceum*. $\wp'$ were calculated from an elliptic curve $[0, 1]$ , expanded to 30-th order, and differentiated.

**Table 7:** *℘*′ values II.

| $\wp'$ | WW *P. pallidum* | WW *D. purpureum* | WW *P. violaceum* |
| --- | --- | --- | --- |
| May |  |  |  |
| June |  |  |  |
| July | 1.162e35 + 9.874e34*I | 1.594e11 - 6.432e11*I | 1.231e52 + 4.372e52*I |
| August | -5.395e28 - 4.181e28*I | -9.400e22 + 1.055e23*I |  |
| September | 4.088e38 + 3.182e38*I | 3.442e47 + 6.318e47*I | 14.97 - 8.365*I |
| October | -5.744e28 - 4.312e28*I |  | -1.223e23 + 1.360e23*I |
| November |  |  |  |
| December |  |  |  |
| January |  |  |  |

WW: Washidu West quadrat; *P. pallidum*: *Polysphondylium pallidum*; *D. purpureum*: *Dictyostelium purpureum*; *P. violaceum*: *Polysphondylium violaceum*. $\wp'$ were calculated from an elliptic curve $[0, 1]$ , expanded to 30-th order, and differentiated.

**Table 8:** Hetero-interaction terms.

| hetero-interaction | WE |  | WW |
| --- | --- | --- | --- |
| Pp-Dp (June) | 0.001160 - 0.09419*I | Pp-Dp (Jul) | 0.01142 - 0.3558*I |
| Pp-Pv (June) | 0.01818 - 0.4494*I | Pp-Pv (Jul) | 0.0008154 - 0.08021*I |
| Dp-Pv (June) | 0.001160 - 0.09419*I | Dp-Pv (Jul) | 0.0008154 - 0.08021*I |
| September | 0.09055 - 0.5885*I | August | 0.09149 - 0.6049*I |
| October | 0.03433 - 0.3021*I | Pp-Dp (Sep) | -2.372e8 + 3.704e8*I |
| November | 0.0003791 - 0.05081*I | Pp-Pv (Sep) | 0.008871 - 0.2633*I |
|  |  | Dp-Pv (Sep) | -1.659e8 - 1.791e9*I |
|  |  | October | -3.728e5 - 3.975e5*I |

WE: Washidu East quadrat; WW: Washidu West quadrat; *P. pallidum*, Pp: *Polysphondylium pallidum*; *D. purpureum*, Dp: *Dictyostelium purpureum*; *P. violaceum*, Pv: *Polysphondylium violaceum*.

For further clarification, regarding *℘* as an elliptic function,

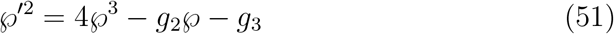

is a normal form without multiple root. Rationals exist, *F* (*℘*(*u*)), *G*(*℘*(*u*)) as Legendre canonical forms of elliptic integrals, such that any elliptic function *f* (*u*) = *F* (*℘*) + *G*(*℘*)*℘*′. Thus a particular state during time procedure *℘*′ can be related to any elliptic function form by a particular pair of Legendre canonical forms. Utilizing Weierstraß *℘* is thus closely related to abstraction of interaction of the states, with a cube of *℘* itself. Setting Ω as a period of *f* (*u*), the canonical form *K*(Ω) ≅ ℂ[*x, y*]/(*y*^2^ − 4*x*^3^ + *g*_2_*x* + *g*_3_), where ℂ[*x, y*] is an integral domain. The ideal thus characterizes the observation phenomena related to *F, G*.

To develop the evaluation, *s* can be regarded as the elliptic function *f* (*u*) via *p*, *l* double periodicity, and a linear plot of *f* (*u*) against *℘*′ shows *F*, *G* values. Basically, due to empirically massive values for *℘*′, *G* ~ 0 and *F* are almost identical to either of the s values selected for calculating the interaction. By this method, one can evaluate which of the interacting partners plays a major role in the interaction. The results are shown in Table 9; in WE, the climax species Pp dominated, while in WW, pioneering species Dp and Pv had significant roles [3]. Note that *F, G* are solutions for corresponding hypergeometric differential equations. Thus *g*_2_, *g*_3_ become apparent during the time development process. *ω* can be calculated by 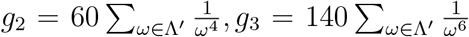. Riemann’s theta relations showed how a (3 + 1)-dimensional system could be rearranged to a 2 + 2 system. Tables 10 & 11 show calculated values for *g*_2_, *g*_3_ in normal form of the elliptic curves.

**Table 9:** *F* values and contributions.

| $F$ | WE | major | | WW | major |
| --- | --- | --- | --- | --- | --- |
| Pp-Dp (June) | 0.7693+8.182*I | Pp | Pp-Dp (Jul) | 0.5752+5.331*I | Dp |
| Pp-Pv (June) | 0.7693+8.182*I | Pp | Pp-Pv (Jul) | 1.262+39.31*I | Pp |
| Dp-Pv (June) | 1.258+31.10*I | Pv | Dp-Pv (Jul) | 0.5752+5.331*I | Dp |
| September | 3.078+14.99*I | Pp | August | 2.879+13.80*I | Dp |
| October | 4.907+38.67*I | Dp | Pp-Dp (Sep) | 1.790+53.12*I | Pp |
| November | 0.7481+7.726*I | Pp | Pp-Pv (Sep) | 0.3186+2.028*I | Pv |
|  |  |  | Dp-Pv (Sep) | 0.3186+2.028*I | Pv |
|  |  |  | October | 2.905+13.93*I | Pv |

WE: Washidu East quadrat; WW: Washidu West quadrat; Pp: *Polysphondylium pallidum*; Dp: *Dictyostelium purpureum*; Pv: *Polysphondylium violaceum*. Major: species that had a major impact on dynamics.

**Table 10:** *g*_2_, *g*_3_ values I.

| Normal form | $g_2$ | $g_3$ | int. | const. | synch. |
| --- | --- | --- | --- | --- | --- |
| Pp-Dp (June) | 3.066e103-2.529e103*I | -7.698e119+1.343e119*I | - | + | anti |
| Pp-Pv (June) | -2.408e65-2.899e65*I | 1.298e81+7.295e81*I | + | - | anti |
| Dp-Pv (June) | 3.066e103-2.529e103 | -3.028e135-1.182e136*I | - | + |  |
| September | -3.651e58-4.368e58*I | -3.373e81-4.044e82*I | + | + | for |
| October | 1.159e75-2.991e73*I | -1.859e110+8.674e109*I | - | + | anti |
| November | 3.747e118-1.663e118*I | -1.630e134-3.473e132*I | - | + |  |

int.: positive or negative effect of an interaction term on $\varphi'$ dynamics; const.: positive or negative effect of a constant on $\varphi'$ dynamics; synch.: coupling between $g_2$ and $g_3$ against the dynamics.

**Table 11:** *g*_2_, *g*_3_ values II.

| Normal form | $g_2$ | $g_3$ | int. | const. | synch. |
| --- | --- | --- | --- | --- | --- |
| Pp-Dp (Jul) | -4.118e70-1.788e71*I | 2.322e82+2.096e81*I | + | - | anti |
| Pp-Pv (Jul) | -1.728e107+2.700e107*I | -6.829e142+7.054e141*I | + | + | for |
| Dp-Pv (Jul) | 1.691e198-1.728e107*I | -3.697e118+1.709e118*I | - | + | anti |
| August | -1.060e58-6.532e57*I | -2.706e79-8.852e80*I | + | + | for |
| Pp-Dp (Sep) | -9.168e77-4.559e78*I | -5.398e116-7.349e116*I | + | + | for |
| Pp-Pv (Sep) | -1.199e78-3.676e78*I | -1.584e79+1.834e78*I | + | + |  |
| Dp-Pv (Sep) | 1.099e77-1.162e77*I | -3.794e77-5.397e77*I | - | + |  |
| October | 1.634e46-5.905e45*I | 4.963e63+3.128e64*I | - | - |  |

int.: positive or negative effect of an interaction term on $\varphi'$ dynamics; const.: positive or negative effect of a constant on $\varphi'$ dynamics; synch.: coupling between $g_2$ and $g_3$ against the dynamics.

## 4 Discussion

Here we move to some more miscellaneous parts associated with eliminating fluctuations. Regarding the utilization of hyperbolic geometry (logarithmic-adic space) and blowing up for resolution of singularity, see our earlier work [5]. From generalized function theories, the idea of cohomology naturally emerges and if we set operator III in terms of cohomology, the *H^p^* = 0(*p* ≥ 1) (p are primes and 1) cohomology and the Kawamata-Viehweg vanishing theorem are fulfilled. This clearly demonstrates that investment in adaptation in the higher order hierarchies diminishes chaotic behavior in the hierarchies. This is because our complex manifold is a Stein manifold (s is a Schwartz distribution). Furthermore, an empirical process is already introduced as “Paddelbewegung” in [1], inspired by Hermann Weyl’s work. Other possible developments for this work include utilizing a Riemann scheme and hypergeometric differential equations or Painlevé VI equations for the hierarchical time-developing model. Consideration of an array of model types would plausibly allow exploration in relation to Galois theory and étale cohomology to interpret the hierarchical structures of natural systems, especially in biological contexts. This thus represents fruitful terrain for future research.

Finally, adopting the Atiyah-Singer index theorem, a twisted (fractal) property, Euler number of ∫_*B*_ *e*(*TB*) is obviously equal to its topological Euler characteristic, χ(*B*) = ∑(−1)^*l*^*l*. Hence, the analytical index of Euler class (Poincaré dual) should be the same. For evaluation of agreement, the Chern class should be (−1)^*l*^*l*. On the other hand, analytically, the Hirzebruch signature (characteristic from species) of *B* is 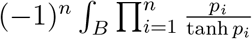, where 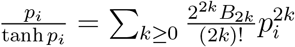. Topologically, this is equivalent to the L genus.

We are thus able to extend the methodology for the “small *s*” metric to characterize dynamical system hierarchy (adaptation and contributions) and interactions, using only abundance data along time development.

## Acknowledgments

This research was primarily funded by the Center of Innovation (COI) Program of Japan Science and Technology Agency. Additional funding was provided by Kyoto University. I extend my gratitude to all the reviewers and colleagues who provided useful information and insights which helped to materially improve this work.

## Notes

#### Summary of Updates

p. 2, middle: and\hat{\mathbb{C}}, `and' deleted; p. 14, middle: 1.001511 -> 1.001571; p. 20, : `$P_0, P_1, P_2$ correspond to $\Re(v), \Re(v'), \Re(v'')$. For $\Re(v), \Re(v'')$, values close to zero represent large contributions, and for $\Re(v')$, large values represent large contributions. The inverses of $\Re(v'), \Re(v'')$ scale for $\Re(v')$' actually mean `$P_0, P_1, P_2$ correspond to $\Re(v'), \Im(v'), \Re(v'')$. For $\Re(v'), \Re(v'')$, values close to zero represent large contributions, and for $\Im(v')$, large values represent large contributions. The inverses of $\Re(v'), \Re(v'')$ scale for $\Im(v')$'. Some other typos in references are corrected.

